# Contrast Subgraphs Allow Comparing Homogeneous and Heterogeneous Networks Derived from Omics Data

**DOI:** 10.1101/2022.07.26.501547

**Authors:** Tommaso Lanciano, Aurora Savino, Francesca Porcu, Davide Cittaro, Francesco Bonchi, Paolo Provero

**Affiliations:** Sapienza University of Rome, 00185 Rome, Italy; Department of Molecular Biotechnology and Health Sciences, Molecular Biotechnology Center, University of Turin, 10126 Turin, Italy; Center for Omics Sciences, San Raffaele Scientific Institute IRCSS, 20132 Milan, Italy; Centai, 10138 Turin, Italy; Eurecat, 08005 Barcelona, Spain; Department of Neurosciences “Rita Levi Montalcini”, University of Turin, 10126 Turin, Italy

**Keywords:** Contrast Subgraphs, Gene Networks, Coexpression Networks, Protein Interaction Networks

## Abstract

Biological networks are often used to describe the relationships between relevant entities, in particular genes and proteins, and are a powerful tool for functional genomics. Many important biological problems can be investigated by comparing biological networks between different conditions, or networks obtained with different techniques. We show that contrast subgraphs, a recently introduced technique to identify the most important structural differences between two networks, provide a versatile tool for comparing gene and protein networks of diverse origin. We show in three concrete examples how contrast subgraphs can provide new insight in functional genomics by extracting the gene/protein modules whose connectivity is most altered between two conditions or experimental techniques.

## Introduction

The development of high-throughput methods in the last few decades has revolutionized biology by allowing the investigation of living systems from a global point of view, thanks to the various omics technologies [16]. The huge amount of data thus produced present new analytical challenges for their interpretation and the extraction of useful and actionable biological information.

An important approach to such analytical task proceeds through the generation, from the high-throughput data, of *biological networks* expressing various types of relationships between the biological entities that have been measured (see [18] for a recent review). In some cases, the results of high-throughput measurements can be directly interpreted as networks, as in the case of protein interaction networks. In other cases, a network structure is built as an analytical tool to facilitate the extraction of biological information, as in the case of coexpression networks in which edges are established between genes showing correlated expression profiles in transcriptomic or proteomic assays. Many analytical tools developed in the context of network science can then be applied to such networks to extract biological information and formulate mechanistic hypotheses.

In many cases of biological interest, the most important questions can be answered not by simply analyzing a single biological system, but by comparing two such systems to extract their fundamental differences. For example, when studying a disease it is necessary to compare the diseased status to the normal one, or different types of disease to each other. Moreover, different omics techniques can produce complementary insights into biological systems, and the investigation of such differences can shed light on the biological features best represented by each technique. When the system of interest has been described in terms of a biological network, techniques for network comparisons become the main tool for these investigations.

The bulk of the methods for network comparison can be categorized into two main classes: methods for the structural comparison of networks and methods for network alignment. Methods in the former class aim to detect global differences between networks in terms of the features considered in network science, such as connectivity distribution, clustering coefficient, assortativity, etc., and do not explicitly identify the individual nodes responsible for such differences. Methods for network alignment are mostly used to identify homologous modules in networks of different origin, and are thus conceived to find similarities, rather than differences, between networks.

Recently, Lanciano et al. [19] proposed the extraction of *contrast subgraphs* as a method to identify the most important structural differences between two networks sharing the same nodes. In essence, contrast subgraphs are sets of nodes whose induced subgraphs are densely connected in one network and sparsely in the other (mathematical definitions and algorithms are found in the Methods). Contrary to most methods for structural comparison, contrast subgraphs are characterized by node identity awareness, *i*.*e*. identify the individual nodes that are responsible for the major differences between the networks. This limits their application to pairs of networks sharing the same nodes, but, on the other hand, allows rich downstream analyses based on domain-specific knowledge on the nodes. For instance, applications in which contrast subgraphs have been employed are social media [33] and neuroscience [19].

Here we apply contrast subgraphs to several comparisons of biological networks derived from high-throughput data, and we demonstrate how meaningful and novel biological information can be extracted from such comparisons. In particular, with respect to existing methods [2], contrast subgraphs exhibit two important advantages. First, the same technique can be used to compare homogeneous networks (that is, obtained from the same high-throughput assay applied to different systems, such as co-expression networks obtained from two different types of cancer) or heterogeneous ones (obtained from different assays, such as protein co-expression and mRNA co-expression networks). Second, the method produces a hierarchically organized list of differentially connected modules that can be interpreted as representing separate biological processes.

## Results

To demonstrate how contrast subgraphs are useful in extracting biological information from the comparison of biological networks we discuss three concrete examples, where the techniques is applied to homogeneous networks (coexpression networks and protein-protein interaction [PPI] networks from different biological conditions) or heterogeneous ones (coexpression networks derived from transcriptomic and proteomic data).

### A. Coexpression networks in two subtypes of breast cancer

Transcriptomic assays have revealed that breast cancer is, from the molecular point of view, a highly heterogeneous disease. The most commonly used transcriptomic-based classification of this disease includes five subtypes (luminal-A, luminal-B, HER-2-enriched, basal-like, and normal-like), where luminal-A and basal-like are considered, respectively, the least and most aggressive subtypes [32]. We used two large repositories of breast cancer gene expression data, namely the TCGA (https://www.cancer.gov/tcga) [23] and METABRIC [6], to build coexpression networks separately for tumors classified as basal-like and as luminal-A. We then extracted the contrast subgraphs from the comparison of the two subtype-specific networks, separately for each dataset.

Figure 1 and Figure 2 (A and B) represent the degree distribution for the first contrast subgraphs showing, as expected, a strong difference between the two subtypes (the genes in the first contrast subgraphs are listed in Suppl. Table 1). Analysing these genes’ enrichment for functional categories, as annotated in the Gene Ontology (GO), we found immune-related processes to be coherently more coexpressed in the basal-like subtype, both in the TCGA and in the METABRIC cohort, while other processes related to tumor microenvironment, such as extracellular matrix organization, are more strongly coexpressed in the luminal-A subtype (Figure 1 and 2 (C and D)). This indicates that the tumor microenvironment, and in particular immune cells and fibroblasts, play a prominent role in differentiating these two molecular subtypes. The full list of enriched GO categories is provided in Suppl. Table 2. Importantly, the results obtained with the two independent breast cancer cohorts show good agreement, with the top differential subgraphs significantly overlapping for both the basal-like and the luminal-A subtypes (Fisher test *p* < 2.2 · 10^−16^), supporting the reliability of the method.

**Fig. 1.**
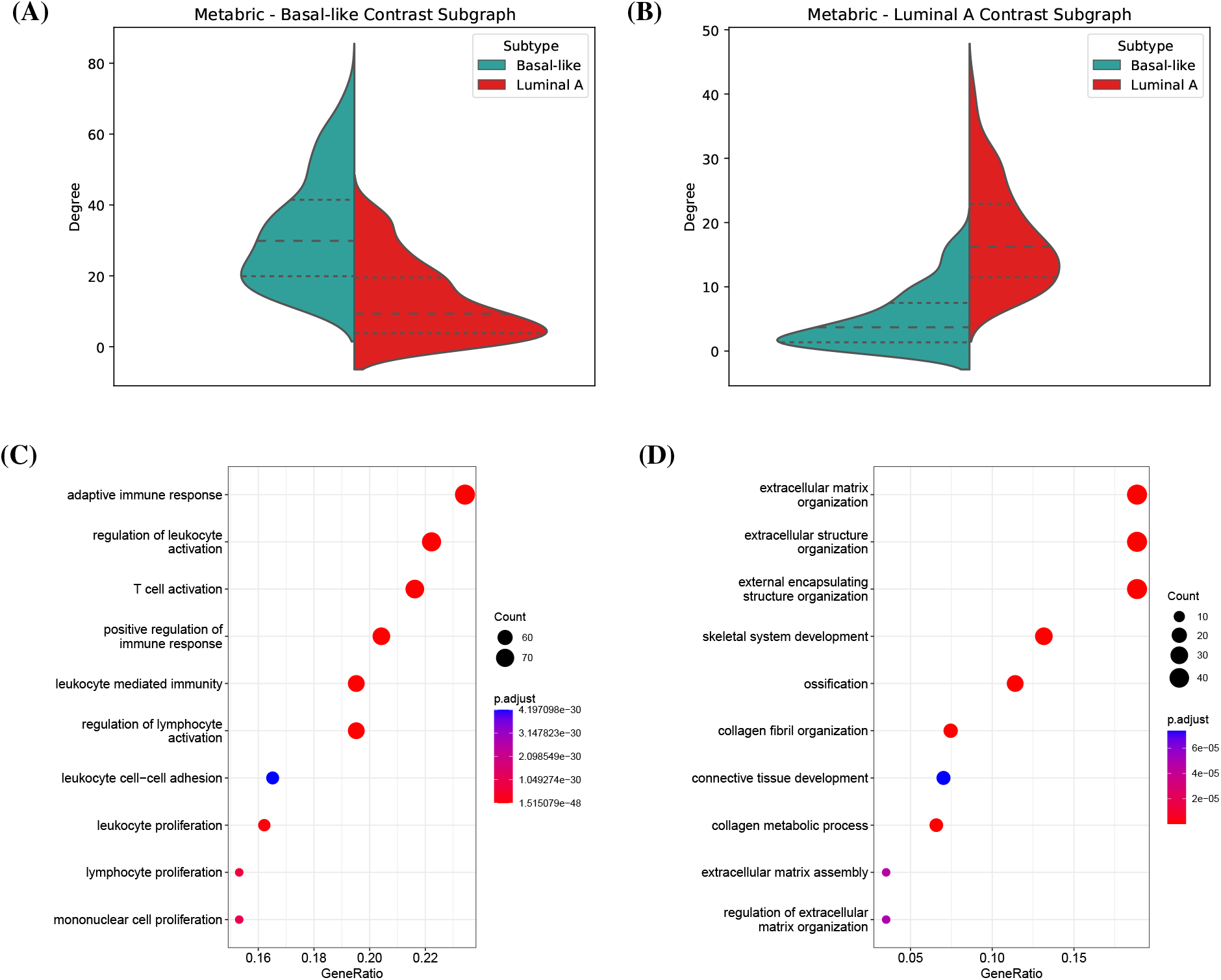
Contrast subgraphs for Metabric. A,B) Degree distributions for both the coexpression networks involved, and quartiles of these distributions. C,D) Dotplots showing the enrichment of each contrast subgraph for Gene Ontology biological processes. The colour gradient indicates the false discovery rate, while the dot size correlates with the number of nodes in the intersection between the contrast subgraph and the functional category. Only the top 10 most significant categories are shown.

**Fig. 2.**
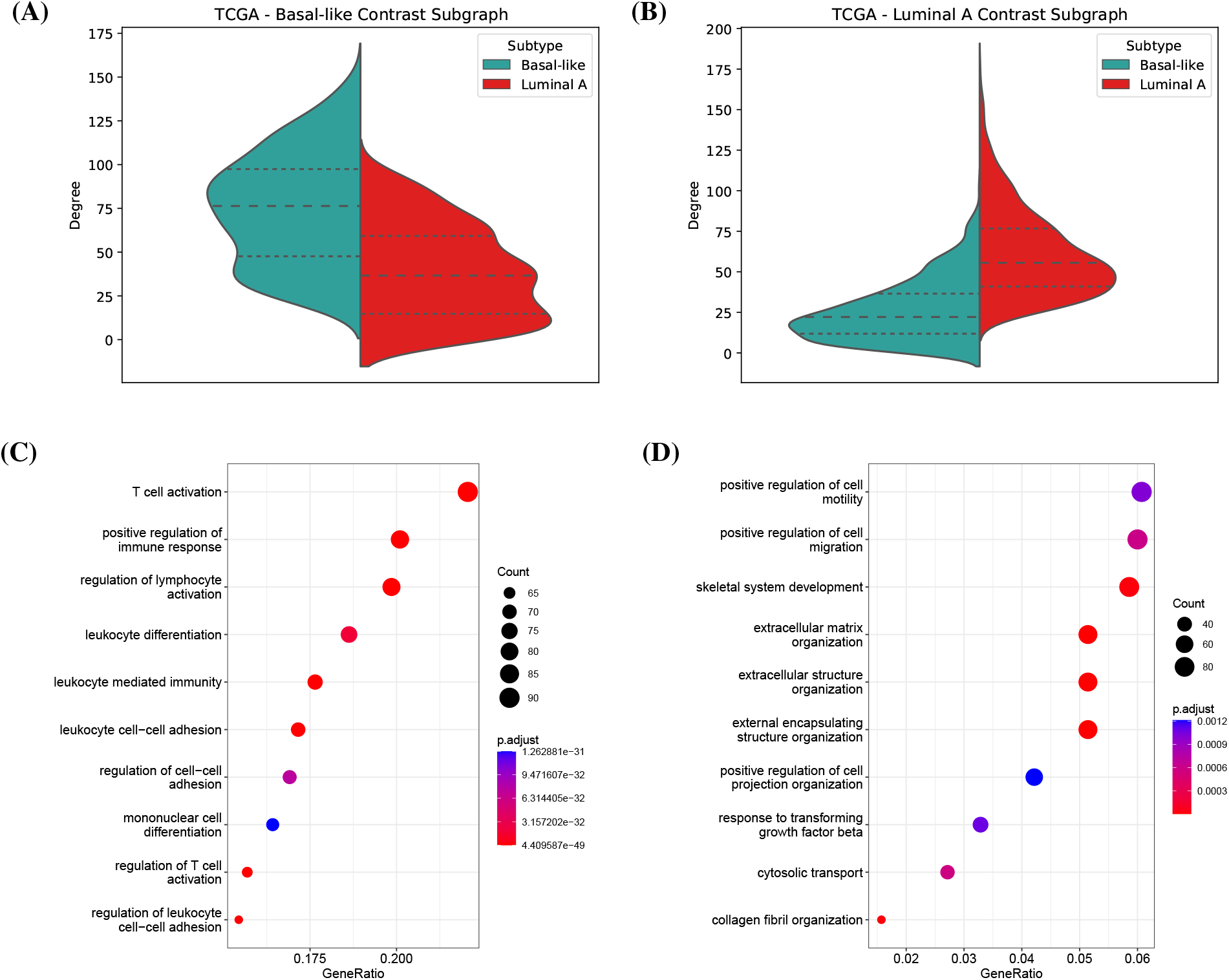
Contrast subgraphs for TCGA. A,B) Degree distributions for both the coexpression networks involved, and quartiles of these distributions. C,D) Dotplots showing the enrichment of each contrast subgraph for Gene Ontology biological processes. The colour gradient indicates the false discovery rate, while the dot size correlates with the number of nodes in the intersection between the contrast subgraph and the functional category. Only the top 10 most significant categories are shown.

### B. Protein vs mRNA coexpression in breast cancer

Coexpression networks are usually built, as we did above, from the results of transcriptomic assays, since these are less expensive than proteomic assays and thus available in large numbers. However, proteins, rather than mRNA molecules, are the predominant components of the molecular machinery performing cellular functions. Moreover, although transcriptomics studies commonly assume mRNA levels to be reliable indicators of corresponding protein levels, transcript and protein expression do not always correlate [20]. Indeed, synthesis and degradation rates of the two types of molecules can be substantially different [35]. Additionally, a wide range of post-transcriptional regulatory mechanisms, among which translational repression by small noncoding RNAs and localization in P bodies, could account for such discrepancies [36]. Therefore it is reasonable to expect that coexpression networks built from protein abundance data could provide information that is complementary to that provided by mRNA-based coexpression networks, and possibly more biologically relevant.

To analyze the differences between mRNA-based and protein-based coexpression networks we used the proteomic data available from CPTAC [22], for a subset of the breast cancer patients included in the TCGA, and built a protein-based coexpression network, which was then compared using contrast subgraphs to the coexpression network obtained from the mRNA data of the same subset of patients.

The subgraphs with the strongest differential coexpression between the proteomic and transcriptomic data (listed in Suppl. Table 3) are enriched for immune categories. Of note, genes more connected at the protein level belong to categories such as “complement activation” and “regulation of humoral immune response”, while genes with functions in the adaptive immunity are over-represented among those with higher transcriptional coexpression (Figure 3, full list in Suppl. Table 4). Moreover, the subgraph more connected at the protein level comprises genes with strikingly low correlation in their mRNA and protein expression (Figure 3, C), indicating that these genes are subject to additional regulatory layers, thus supporting their discrepant mRNA and protein coexpression. This observation is in line with the complement cascade being mostly regulated through proteolytic activity, and indicates that subgraphs more connected at the proteome level could better represent functional coupling of processes regulated at the post-translational level.

**Fig. 3.**
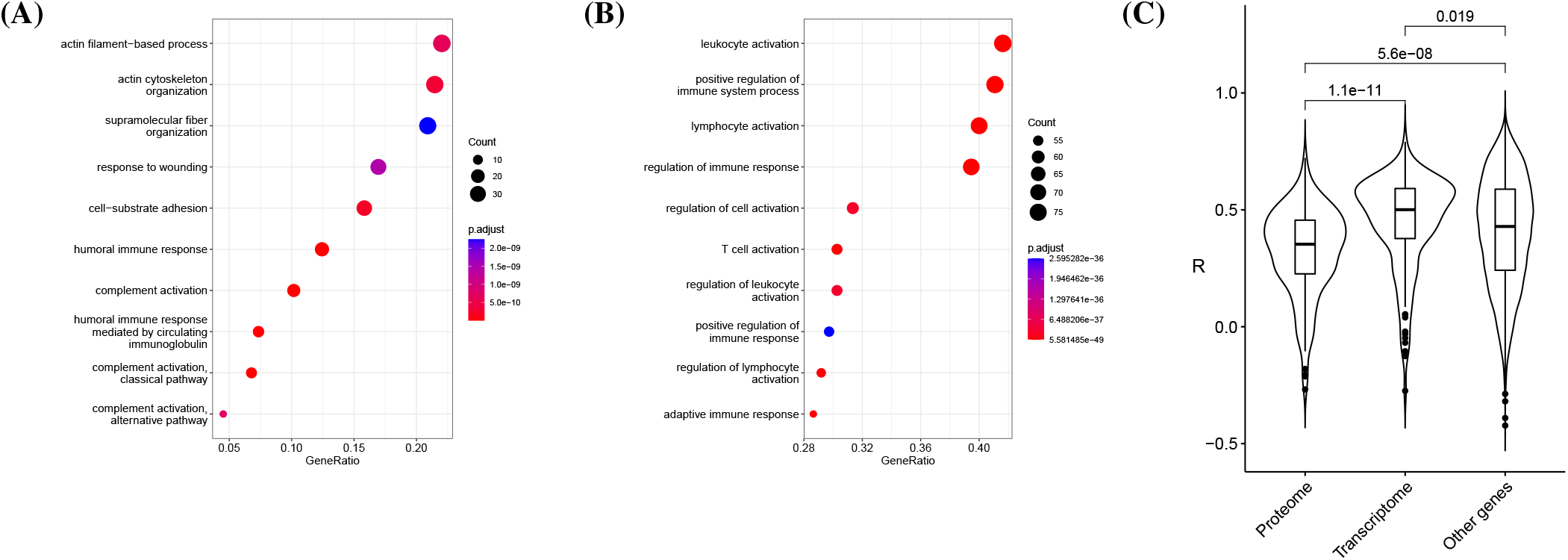
A, B) Dotplots showing the enrichment of each contrast subgraph for functional categories. The colour gradient indicates false discovery rate, while dot size correlates with the number of nodes in the intersection between the contrast subgraph and the functional category. Only the top 10 most significant categories are shown. C) Violin plot showing Pearson’s correlation between transcriptomic and proteomic levels for genes in the top differential subgraph most connected at the proteome or at the transcriptome level, compared with genes not belonging to any of the two subgraphs.

### C. Protein interaction networks in human cell lines

The analysis of PPI networks can provide functional information complementary to that provided by transcriptomics. In particular, the comparison of such networks derived from different cell types or tissues can indicate those interactions that are specific to a biological context. We thus considered experimentally determined PPI networks in three human cell lines (HUVEC, HEK293T, and JURKAT) [17], and extracted the contrast subgraphs for each of the 6 possible comparisons. Each contrast subgraph thus contains proteins with higher density of interactions in one cell line compared with the other.

It is important to verify that the contrast subgraphs thus extracted do not simply contain proteins that are differentially expressed when comparing the three cell lines. We compared the first contrast subgraphs obtained for each comparison with lists of upregulated genes in the same comparison derived from the RNA-seq data of the Human Protein Atlas [34]. While the proteins contained in the first contrast subgraph in HUVEC and, to a lesser extent, JURKAT cells did indeed significantly overlap the corresponding up-regulated genes, such enrichment was not detected in HEK293T-specific contrast subgraphs, which we can confidently consider as representing true cell type-specific interaction modules.

We thus analyzed in more depth the first contrast subgraphs obtained from the comparison of the HEK293T PPI network with those obtained from the other two cell lines. Remarkably, the top contrast subgraphs obtained from the two comparisons were identical, and contained 143 proteins (Suppl. Table 5). Gene Ontology enrichment analysis of these proteins revealed 160 enriched biological processes (Suppl. Table 6), including many terms related to translation and ribosome biogenesis on one hand, and many related to apoptosis and the TP53 pathway on the other. Figure 4 shows the proteins annotated “ribosome biogenesis” and “signal transduction by p53 class mediator” and their interactions in the HEK293T and HUVEC cell lines. These results suggest that HEK293T cells are particularly suitable for the investigation of the deep relationship between TP53 and the ribosome [7]. Indeed these cells have been used in the original experimental investigation of this relationship [30].

**Fig. 4.**
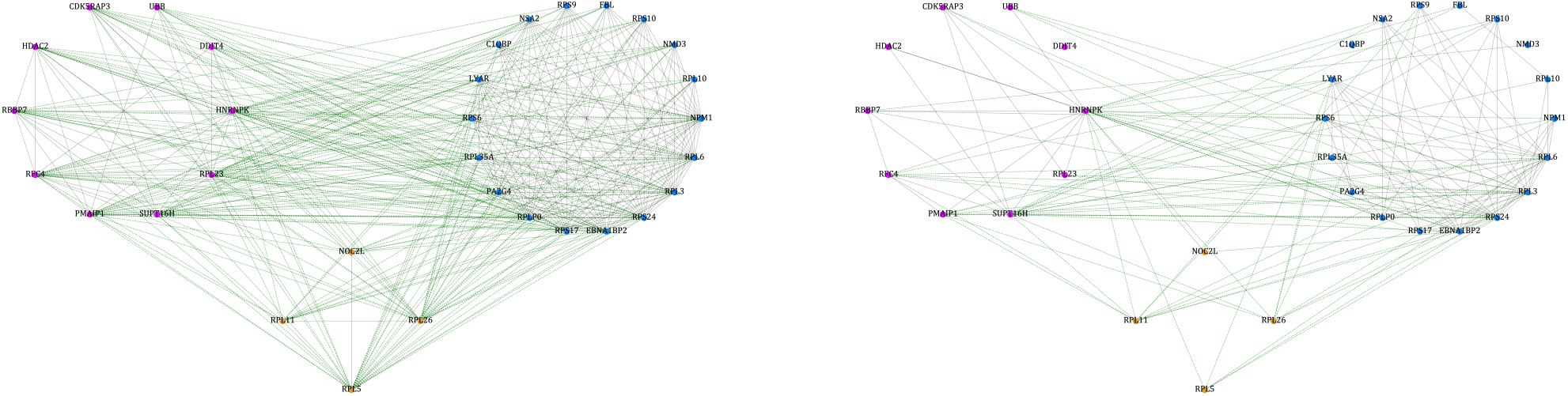
Proteins involved in “ribosome biogenesis” (blue), “signal transduction by p53 class mediator” (purple) and in both processes (yellow) together with their interactions in HEK293T (left) and HUVEC cells (right). The green edges join proteins involved in the two biological processes. The abundance of green edges in the left panel illustrates that the many interactions between these two processes are specific of the HEK293T cellular context.

These results show that contrast subgraphs can be used to identify cell-type specific modules of interacting proteins, thus facilitating the choice of the cells to be used for experimental assays.

## Methods

### A. Extraction of contrast subgraphs

Mining contrastive structures from networks has started recently to gain attention in the scientific literature. In this work we leverage this recent literature to provide a first proposal of mining *contrast subgraphs* in the biological domain. Given two (potentially weighted) networks *A* = (*V, e*_*A*_(*V*)) and *B* = (*V, e*_*B*_(*V*)) defined over the same set of nodes *V*, we define a contrast subgraph as a set of nodes that is densely connected in one of the networks, and sparse in the other. In order to quantify this property, different definitions of *contrast* have been provided in the literature.

Lanciano et al. [19] define the contrast subgraph as the set of nodes *S* ⊆ *V* that maximizes the function 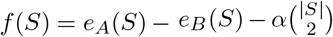, where *e*_*A*_(*S*) and *e*_*B*_(*S*) are the number of edges (or the sum of edges’ weight in case of a weighted network) in the subgraph induced by *S* in the networks *A* and *B*, respectively, and *α* is an input scalar. This definition aims at identifying a set of nodes, whose induced subgraph is dense in *A* and sparse in *B*. The regularization term 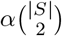, governed by the parameter *α*, can be used to tune the target size of *S*: in fact, all the edges whose weight is smaller than *α* giving a negative contribution to the objective function, thus preventing larger solutions. To maximize this function, the authors map their problem to an instance of the Generalized Optimal Quasi Clique problem proposed by Cadena et al. [3]. Their algorithm is based on an Semi-Definite Programming optimization problem, that makes it practical only for smaller instances of networks, e.g. brain networks.

A variant formulation for this problem, by Yang et al. [37], aims at maximizing 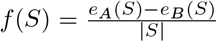: they show that their problem is NP-hard and proposed a simple heuristic. It is worth observing that their problem corresponds to the classic *Densest Subgraph Problem* (DSP) [12] on a weighted network, where the weight of an edge is given by *e*_*A*_(*S*) -*e*_*B*_(*S*). DSP is one the most important primitives in graph mining, that has been studied extensively in literature for its many potential applications. Given a graph, it aims at finding the subgraph with maximum average degree, i.e., 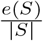. When the graph is unweighted or positively weighted, DSP can be solved exactly in polynomial time by means of an inefficient max-flow based algorithm [12]. An efficient 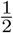 -approximation of the exact solution, can be obtained by a greedy “peeling” algorithm that at every iteration removes the nodes with the current minimum degree, and among all intermediate subgraphs produced by this process, in the end it returns the one maximizing the objective function [1, 4].

Unfortunately, when the graph has weights that can be negative, as in our case, these algorithmic results do not carry on (indeed, the contrast subgraph problem by Yang et al. [37] is NP-hard).

Tsourakakis et al. [33] recently analyzed the performance of the greedy peeling for the *Densest Subgraph with Negative Weights* (DSNW) problem. Let *deg*^+^(*v*) be the positive degree of node *v*, i.e., the sum of the weights of its positive edges, and *deg*^−^(*v*) its negative degree. They provide the following lower bound on the solution’s quality: 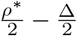, where *ρ*^*^ is the optimum value of the DSP problem, and 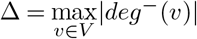.

In order to improve such result, they propose a variant of the greedy peeling (Algorithm 1), introducing a parameter *C* that governs the importance of *deg*^+^(*v*) in order to avoid the bad instances for which the greedy peeling could fail. It is sufficient to tune this parameter and run several times Algorithm 1 to obtain a better result without a significant increase in computing time. Given its efficiency and scalability, and the fact that it has a certified lower bound on the quality of the quality of the solution provided, in our experiments we employ Algorithm 1 to mine the contrast subgraphs of coexpression and PPI networks.

In the literature reviewed above, the contrast subgraph is the one subgraph maximizing the contrastive objective function. However, although according to our Algorithm 1 the extraction is limited to the subgraph that maximizes the contrast function, a straightforward heuristic to mine the top-*k* non-overlapping contrast subgraphs can be easily implemented, by simply iterating Algorithm 1 for *k* times, removing from the graph at each iteration the nodes obtained in output.

### B. Construction of coexpression networks

Normalized (FPKM) breast cancer data from the TCGA project and corresponding clinical annotations were obtained through TCGA biolinks [5], and METABRIC gene expression data and metadata were obtained from www.synapse.org (Synapse ID: syn1688369) [6]. Probe names were converted in Gene Symbols and, for gene symbols corresponding to multiple probes, the most expressed probe across all samples was considered. ENSEMBL IDs were converted into gene symbols using biomaRt [9]. In TCGA, genes with FPKM < 1 in more than 50 samples were filtered out, and data were log transformed using an offset of 1. Proteomic CPTAC data were downloaded from the original publication [22]. Adjacencies were computed using Spearman’s correlation between gene or protein expression, then transformed as follows to generate a signed network (0.5 · (1 + *ρ*))^12^, where *ρ* indicates Spearman’s correlation. Functional enrichment was performed using clusterProfiler [39], considering only categories with an adjusted p-value less than 0.05.

#### Algorithm 1 Heuristic Peeling [33]

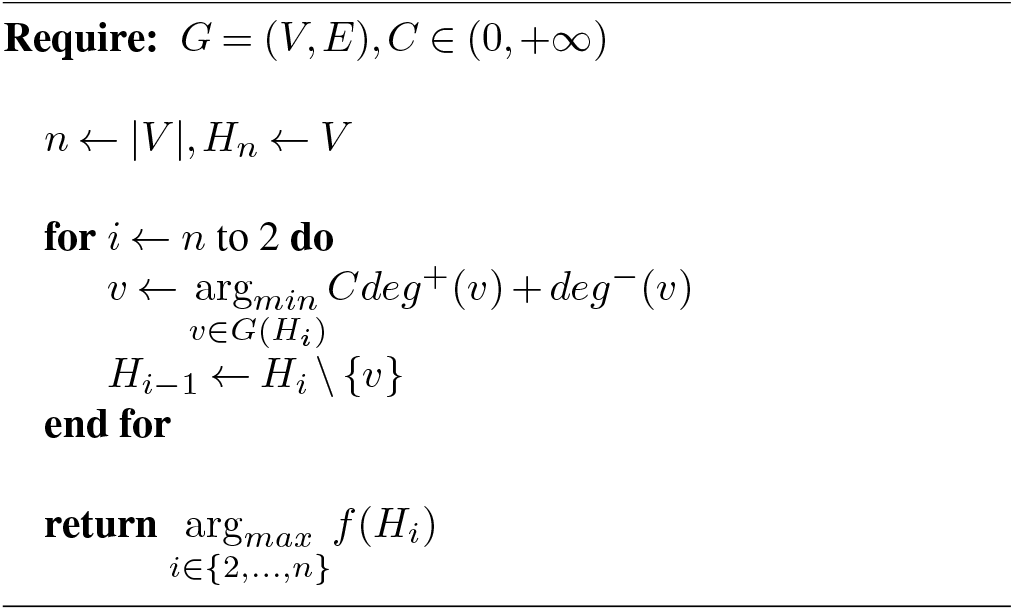

### C. Analysis of PPI constrast subgraphs

PPI networks obtained in HEK293T, HUVEC, and JURKAT cells were obtained from the Supplementary Material of the original publication [17]. PPI networks were filtered to include only proteins described in all cell lines. RNA-seq data for the same cell lines were obtained form the Human Protein At-las [34]. As these data do not contain replicates, lists of genes upregulated in each comparison were obtained by requiring |*logF C*| *>* 1 on the normalized TPM values after logarithmic transformation with unit pseudocount.

## Discussion

Gene networks have proved to be a valuable tool to understand some general principles governing biological systems, revealing a modular organization of gene interactions [25], at least partly linked to shared function [10]. Comparing molecular networks across different contexts is nevertheless essential to explore the biology of dynamic systems, with gene and protein interactions changing over time or upon perturbations, such as disease or environmental stresses.

Many of the methods that have been developed for the comparison of biological network focus on structural properties and often lack node identity awareness; other methods focus on network alignment, and aim at finding commonalities, rather than differences, between networks. Here we have shown that contrast subgraphs can provide a versatile tool to identify the modules with the strongest difference in connectivity between two networks. The method can be applied to networks of different biological and technical origin, and its node identity awareness allows downstream biological analyses providing insight on the biological processes affected by differential connectivity. It should be noted that for the specific case of co-expression networks, several comparative methods that do retain node identity awareness have been developed (reviewed in [2, 29]). However most of these methods are specifically targeted to correlation-based networks and are not immediately applicable to other biological networks such as PPI networks.

In this work, we applied contrast subgraphs to three pairs of biological networks to illustrate their usefulness, especially when followed by downstream functional enrichment analysis of the differential modules identified.

Breast cancer (BC) is one of the leading causes of mortality in women worldwide, for which no general efficacious treatment is available due to disease heterogeneity. BC is usually classified according to gene expression profiling of the tumor (PAM 50 assay) allowing the classification in five molecular subtypes correlated with prognosis and response to treatments: luminal-A, luminal-B, basal-like, HER2-positive and normal-like [24,31]. In particular, basal-like subtype does not respond to targeted treatments such as hormonal blockage or Herceptin, and shows poor outcome.

We analyzed the different organization of co-expression networks between the aggressive basal-like and the relatively slowly growing luminal-A subtypes. We find that the top differentially connected subgraphs comprise genes enriched for microenvironment-related functions in two independent cohorts with associated transcriptome datasets (METABRIC [6] and TCGA [23]). On one side, the subgraph with stronger connection in the basal-like subtype is enriched for immune functions, while, on the other side, the genes more connected in the luminal-A subtype are enriched for categories such as “Extracellular Matrix”, indicative of microenvironmental regulation of structural components of the extracellular milieu. Indeed, tumor cells are surrounded by a varied ensemble of mutually-interacting cell types, comprising immune cells, stromal cells, and blood vessels, amongst the most frequent cell types. These cells can either restrain tumor growth or support cancer cells providing metabolites, growth factors and reshaping the extracellular matrix. Overall, nontumoral cells surrounding the tumor epithelium have been demonstrated to change their expression profiles [28] and to impact not only on tumor growth, but also on disease progression and metastasis, and on drug resistance [27, 38, 40]. In particular, the immune system plays a fundamental role in cancer progression: at tumor onset, cytotoxic immune cells recognize and kill tumor cells, driving the evolution of less immunogenic cancer cells able to evade immune detection [13]. Paradoxically, immune cells such as antiinflammatory M2 macrophages can have pro-tumoral effects [21] and their distribution and composition changes with tumorigenesis [11, 28]. For these reasons, immune cells are currently being investigated as potential therapeutic targets [8, 14]. Interestingly, we previously reported that differentially coexpressed networks between normal and tumor tissues are often enriched for immune-related categories [29], thus confirming that the contrast subgraph method reliably retrieves robust and biologically informative sets of genes.

As a second application context, we employed our differential subgraph retrieval to compare biological networks derived from different kinds of molecular data: transcriptomics and proteomics. The possibility of comparing networks from different data types is indeed a strength of our method. We made use of the large breast cancer TCGA cohort of primary tumors, which have been profiled both through RNA-seq and mass spectrometry [22], and defined gene modules whose connectivity can be revealed only at the proteomic level, likely due to post-transcriptional regulatory mechanisms influencing protein translation and degradation. Intriguingly, the two top differential modules show a significant difference in their transcript-protein agreement, supporting the hypothesis of intervening post-transcriptional mechanisms being involved in the proteome subgraph regulation. Again, these top differential subgraphs are enriched for immune-related categories. Interestingly, adaptive immune system genes are more connected at the transcriptional level, while innate immune system genes are more connected at the protein level. This difference could be interpreted as the innate immune system being poised for a rapid activation in the presence of a stimulus in the form of a pathogen or of a tumor cell, hence relying on a fast and coordinated translation of readily available transcripts. The adaptive immune response, on the other hand, acts more slowly, requiring days or even weeks to become established. Therefore, transcription is not a limiting factor in its response, making transcriptionally-regulated modules easily detectable. Indeed, the proteomic subgraph is significantly enriched for the complement cascade, which is extensively regulated through the activity proteolytic enzymes ([26], [15]).

In the third example we showed that the comparison of PPI networks obtained from different human cell lines can reveal how proteins involved in different biological processes can have context-dependent interaction patterns. Importantly, such differences were not apparent from differential expression analysis of the same cell lines. These results suggest, in particular, that contrast subgraphs can be useful in selecting the cellular contexts most suitable for the experimental analysis of the interaction and mutual dependence of different biological processes.

## Conclusion

Contrast subgraphs are a promising and versatile method to identify the most relevant differences between biological networks while preserving node-identity awareness, thus allowing the translation of such information into biological insight.

## ACKNOWLEDGEMENTS

The results of this analysis are in whole or part based upon data generated by the TCGA Research Network: https://www.cancer.gov/tcga, accessed on 27 January 2021; and data generated by the Clinical Proteomic Tumor Analysis Consortium (NCI/NIH).

## Supplementary Material

Supplementary material is made available at:

https://zenodo.org/record/6802221

Across the supplementary tables, the differential subgraphs have been named according to the following scheme:

- **METABRIC basal**: genes more connected in the basal than in the luminal A subtype in the METABRIC dataset
- **METABRIC lumA**: genes more connected in the luminal A than in the basal subtype in the METABRIC dataset;
- **TCGA basal**: genes more connected in the basal than in the luminal A subtype in the TCGA dataset;
- **TCGA lumA**: genes more connected in the luminal A than in the basal subtype in the TCGA dataset;
- **Transcriptome**: genes more connected at the transcript level than at the protein level;
- **Proteome**: genes more connected at the protein level than at the transcript level.

### Supplementary Tables

- **Supplementary Table 1**. Genes in the first differential subgraphs comparing breast cancer molecular subtypes
- **Supplementary Table 2**. Gene Ontology categories significantly enriched in the corresponding differential subgraph comparing breast cancer molecular subtypes
- **Supplementary Table 3**. Genes in the first differential subgraphs comparing transcriptional and protein co-expression networks.
- **Supplementary Table 4**. Gene Ontology categories significantly enriched in the corresponding differential subgraph comparing transcriptional and protein co-expression networks.
- **Supplementary Table 5**. Genes in the first differential subgraph comparing HEK and Jurkat or HUVEC cell lines
- **Supplementary Table 6**. Gene Ontology categories significantly enriched in the differential subgraph comparing HEK and Jurkat or HUVEC cell lines.

## Software availability

A software package to compute constrast subgraphs is available at

https://github.com/tlancian/bio_cs

